# Intracellular glycosyl hydrolase PslG shapes bacterial cell fate, signaling, and the biofilm development of *Pseudomonas aeruginosa*

**DOI:** 10.1101/2021.08.17.456744

**Authors:** Jingchao Zhang, Huijun Wu, Di Wang, Chenxi Zhang, Kun Zhao, Luyan Z. Ma

## Abstract

Biofilm formation is one of most important causes leading to persistent infections. Exopolysaccharides are usually a main component of biofilm matrix. Genes encoding for glycosyl hydrolases are often found in gene clusters that are involved in the exopolysaccharide synthesis. It remains elusive about the functions of intracellular glycosyl hydrolase and why a polysaccharide synthesis gene cluster requires a glycosyl hydrolase. Here we systematically studied the role of intracellular PslG, a glycosyl hydrolase that is co-transcribed with 15 *psl* genes, which is responsible for the synthesis of exopolysaccharide Psl (ePsl), a key biofilm matrix polysaccharide in opportunistic pathogen *Pseudomonas aeruginosa*. We showed that lacking of PslG in this opportunistic pathogen alters the signaling function and structure of ePsl, changes the relative level of cyclic-di-GMP within daughter cells during cell division and shapes the localization of ePsl on bacterial periphery, thus results in long chains of bacterial cells, fast-forming and compact biofilm microcolonies. Our results reveal the important roles of intracellular PslG on the cell fate and biofilm development.

## Introduction

Structured, surfaced-associated communities of microorganism known as biofilms are important life forms of bacteria prevailing in nature, industrial, and clinical settings (Costerton *et al*., 1995; Stoodley *et al*., 2002). In general, biofilm development involves four specific stages: attachment, microcolony formation, matured microcolonies, and dispersal. Bacteria within biofilms are embedded in extracellular matrix that protect bacterial cells from antibiotics, host defenses, and environmental stresses. Even though the components of biofilm matrix differ from species to species, it generally composes of exopolysaccharides, proteins, and nucleic acids (Stoodley *et al*., 2002; Flemming and Wingender, 2010). Exopolysaccharides are critical biofilm matrix components for many bacteria, which often promote attachment to surfaces and other cells, act as a scaffold to help maintain biofilm structure, and provide protection (Stewart *et al*., 2001; Haussler and Parsek, 2010) . Gene encoding for glycosyl hydrolase is often found in gene clusters that are involved in the synthesis of exopolysaccharide (Franklin *et al*., 2011). Beyond of exopolysaccharide degradation, little is known about whether these genes affect bacterial physiology and biofilm development.

*Pseudomonas aeruginosa* is an opportunistic human pathogen that can cause life-threatening infections in cystic fibrosis patients and immune-compromised individuals (Govan and Deretic, 1996; Lyczak *et al*., 2000; Ramsey and Wozniak, 2005). In *P. aeruginosa* PAO1, <u>e</u>xopolysaccharide Psl (ePsl) is a primary scaffold matrix component that can form a fibre-like matrix to enmesh bacteria within a biofilm (Colvin *et al*., 2012; Ma *et al*., 2009; Wang *et al*., 2013). ePsl has shown multiple functions in the biofilm formation of PAO1. For example, ePsl can act as a “molecular glue” to promote bacteria cell-cell and cell-surface interactions (Ma *et al*., 2006; Ma *et al*., 2009). ePsl trails on a surface guide bacteria exploration and microcolony formation (Zhao *et al*., 2013). Moreover, ePsl can also work as a barrier to protect bacteria from antibiotics and phagocytic cells (Mishra *et al*., 2012; Billings *et al*., 2013; Tseng *et al*., 2013). Interestingly, ePsl can function as a signal to stimulate biofilm formation through affecting intracellular signal molecule cyclic-di-GMP (c-di-GMP) (Irie *et al*., 2012).

ePsl is synthesized by *psl* operon, containing 15 co-transcribed genes (*pslABCDEFGHIJKLMNO*) (Byrd *et al*., 2009). PslG protein encoded by *pslG* gene is a glycosyl hydrolase. PslG has been shown to degrade ePsl *in vitro* or within biofilm matrix and hence inhibit biofilm formation at a nanomolar concentration (Yu *et al*., 2015; Baker *et al*., 2015; Zhao *et al*., 2018). PslG was first thought to be essential for Psl synthesis, since deletion of *pslG* gene led to a loss of ePsl production (Byrd *et al*., 2009). A later work found that deletion of *pslG* in the earlier work might have affected the gene expression downstream of *pslG* and thus resulted in the loss of ePsl production, and absence of *pslG* itself did not result in a complete loss of ePsl production *per se*, but led to a less production of ePsl and reduced bacterial initial attachment compared with PAO1 (Wu *et al*., 2019). In addition, the glycoside hydrolytic activity of PslG has been shown to play a role in the biosynthesis of ePsl (Wu *et al*., 2019).

C-di-GMP is an important second messenger controlling a wide range of cellular processes in many bacteria, such as motility, cell differentiation, biofilm formation and production of virulence factors (Romling *et al*., 2013). Reports have shown that c-di-GMP is asymmetrically distributed among daughter cells upon bacterial cell division and the asymmetric divisions on surfaces produces specialized cell types, a spreader for dissemination and a striker for local tissue damage (Christen *et al*., 2010; Laventie *et al*., 2019). It has not been investigated whether an intracellular glycoside hydrolase would affect the c-di-GMP level.

In this work, aiming to study the effect of PslG on the cell fate and biofilm development of *P. aeruginosa* at the single cell level, we systematically studied the *pslG* knock-out mutants by employing a high-throughput bacterial tracking technique (Zhao *et al*., 2013). The morphology and motility behavior of bacterial cells in the course of biofilm development were analyzed at the single cell level. Using p*cdrA*::*gfp* reporter, the c-di-GMP level of each cell was also monitored. Microscopically, the attachment behavior of cells on the microtiter surfaces and the pellicles formed at the air-liquid interface were also characterized. Our data suggest that lacking of *pslG* impacts cell morphology, the signaling function of ePsl, the c-di-GMP distribution and bacteria distribution within a biofilm. Based on our results together with those in literature, a model is proposed to understand the role of *pslG* in the biofilm development.

## Results

### Increasing ePsl production cannot recover the defects of Δ*pslG* on bacterial attachment

Our previous study showed that the *pslG* deletion mutant (was named as Δ*pslG*2 by Wu et al., hereafter termed as Δ*pslG*) decreased the production of ePsl and bacterial initial attachment on the microtiter surface (Wu *et al*., 2019). To know whether the attachment defect of Δ*pslG* mutant is due to ePsl reduction, we constructed a Δ*pslG* mutant in WFPA801 background, named as WFPA801Δ*pslG.* WFPA801 is a PAO1-derived Psl-inducible strain, in which the promoter of *psl* operon has been replaced by P_BAD_ promoter (Ma *et al*., 2006). Thus, the ePsl production of WFPA801 depends on the concentration of arabinose (the inducer), which is the same for WFPA801Δ*pslG,* a Psl-inducible Δ*pslG* mutant strain. As shown in Figure 1A, WFPA801Δ*pslG* produced similar amount of ePsl as that of PAO1 when induced with 0.5% arabinose. However, WFPA801Δ*pslG* was not able to form a ring at the air-liquid interface as that seen in either PAO1 or WFPA801 under 0.5% arabinose induction (Figure 1A). These results indicate that ePsl reduction is not the reason for the decrease of attachment of *pslG* deletion mutants.

**Figure 1.**
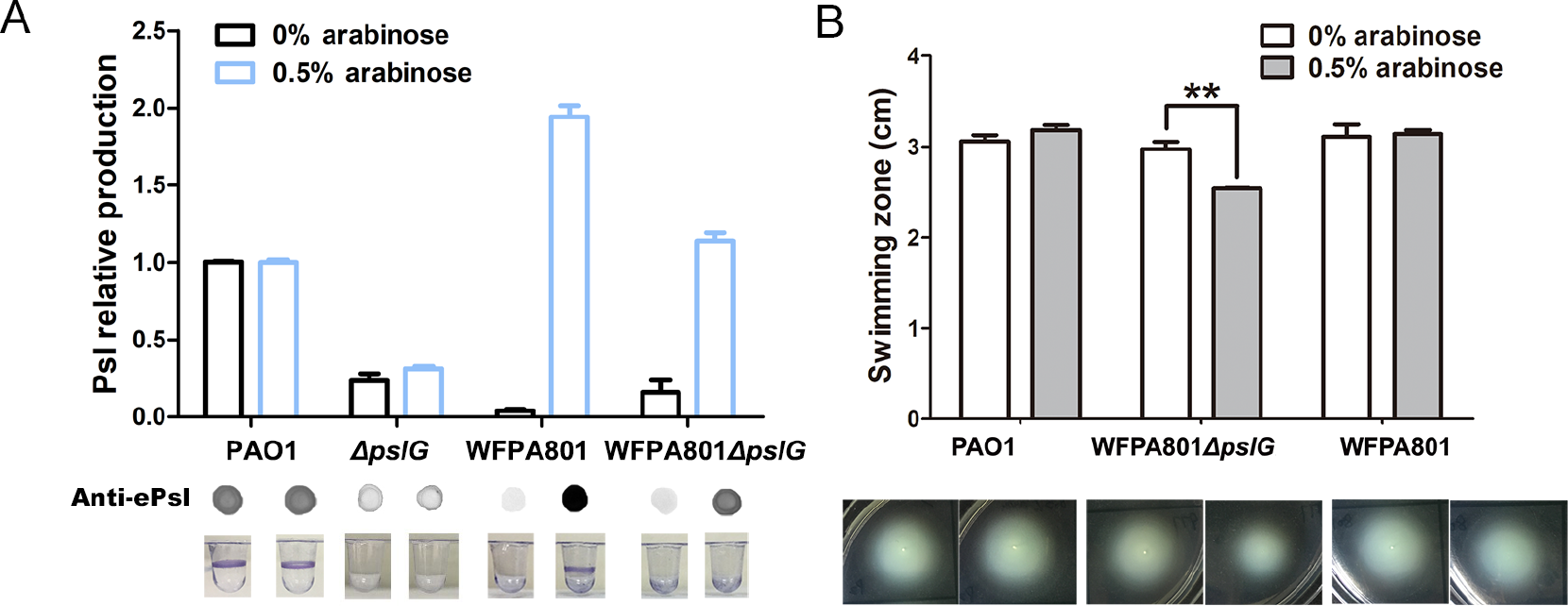
Loss of PslG affects bacterial initial attachment and swimming motility. A: The attachment and Psl production of PAO1, Δ*pslG*, WFPA801, and WFPA801Δ*pslG* inducing with 0% or 0.5% arabinose. The corresponding tubes posted CV staining in attachment assay were shown under each column. B: The swimming motility of PAO1, WFPA801Δ*pslG*, and WFPA801 inducing with 0% or 0.5% arabinose. The corresponding image of swimming zone was shown under each column. Statistical significances were measured using student’s t-test (**, p < 0. 01 ) . **Figure 1-source data -Figure 1A and 1B source data.** **Figure 1-Figure supplement 1. The influence of Δ*pslG* on T4P-driven twitching motility.** A: Tracking T4P-driven bacterial walking and crawling on the glass surfaces of flow-cell systems. Shown are the fractions of cells that crawl only, walk only and both crawl and walk for WFPA801Δ*pslG* and PAO1. B: Speed of cell twitching for WFPA801Δ*pslG* and PAO1. The number of analyzed cells is n=496 for PAO1 and n=576 for WFPA801Δ*pslG*. C: The effect of Δ*pslG* on the twitching zone.

Flagella and Type IV pili (T4P) are also important for initial attachment of *P. aeruginosa* (O’Toole and Kolter, 1998; Bruzaud *et al*., 2015). We then tested flagellum-driven swimming motility and T4P-mediated twitching motility of WFPA801Δ*pslG* strain to evaluate the function of flagella and T4P. Without arabinose induction (the *psl* operon is not transcribed and no ePsl is produced in WFPA801-derived strains, Figure 1A), WFPA801Δ*pslG* strain exhibited similar swimming ability and twitching motility as did PAO1 and WFPA801 (Figure 1B and Figure 1-Figure supplement 1). However, with 0.5% arabinose induction, WFPA801Δ*pslG* strain (having wild type level of ePsl production, Figure 1A) showed reduced swimming zone compared to that of either PAO1 or WFPA801 (Figure 1B), indicating that the induction of ePsl production in WFPA801Δ*pslG* attenuated its swimming ability. However, T4P-mediated twitching motility was not affected (Figure 1-Figure supplement 1C). These results suggested that the defect of Δ*pslG* on initial attachment was a result of multiple contributors.

### **Δ***pslG* impacts the bacterial distribution and maximum thicknesses of pellicles

We then investigated the effect of Δ*pslG* on the air-liquid interface biofilms, termed as pellicles, by using confocal laser scanning microscopy. The total pellicle biomass of Δ*pslG* is similar to that of PAO1 after 24h growth, although Δ*pslG* produced much less ePsl and had defect on initial attachment (Figure 1 and Figure 2A, C). However, Δ*pslG* has significant higher maximum thickness than that of PAO1 (Figure 2B). In addition, there is less bacteria in each section image of Δ*pslG* pellicles compared to that of PAO1 (Figure 2D, left and middle panel). The ePsl matrix in Δ*pslG* pellicles shows weaker fluorescent intensities than that of PAO1 (Figure 2D, middle and right panels), which is consistent with their corresponding ePsl production. In spite of that, the fibre-like ePsl can be detected in the pellicles of Δ*pslG*, which have a radial pattern as previously described for PAO1 pellicles (Figure 2D, middle panels) (Wang *et al*., 2013). These results suggest that the *pslG* deletion might impact bacterial distribution within biofilms.

**Figure 2.**
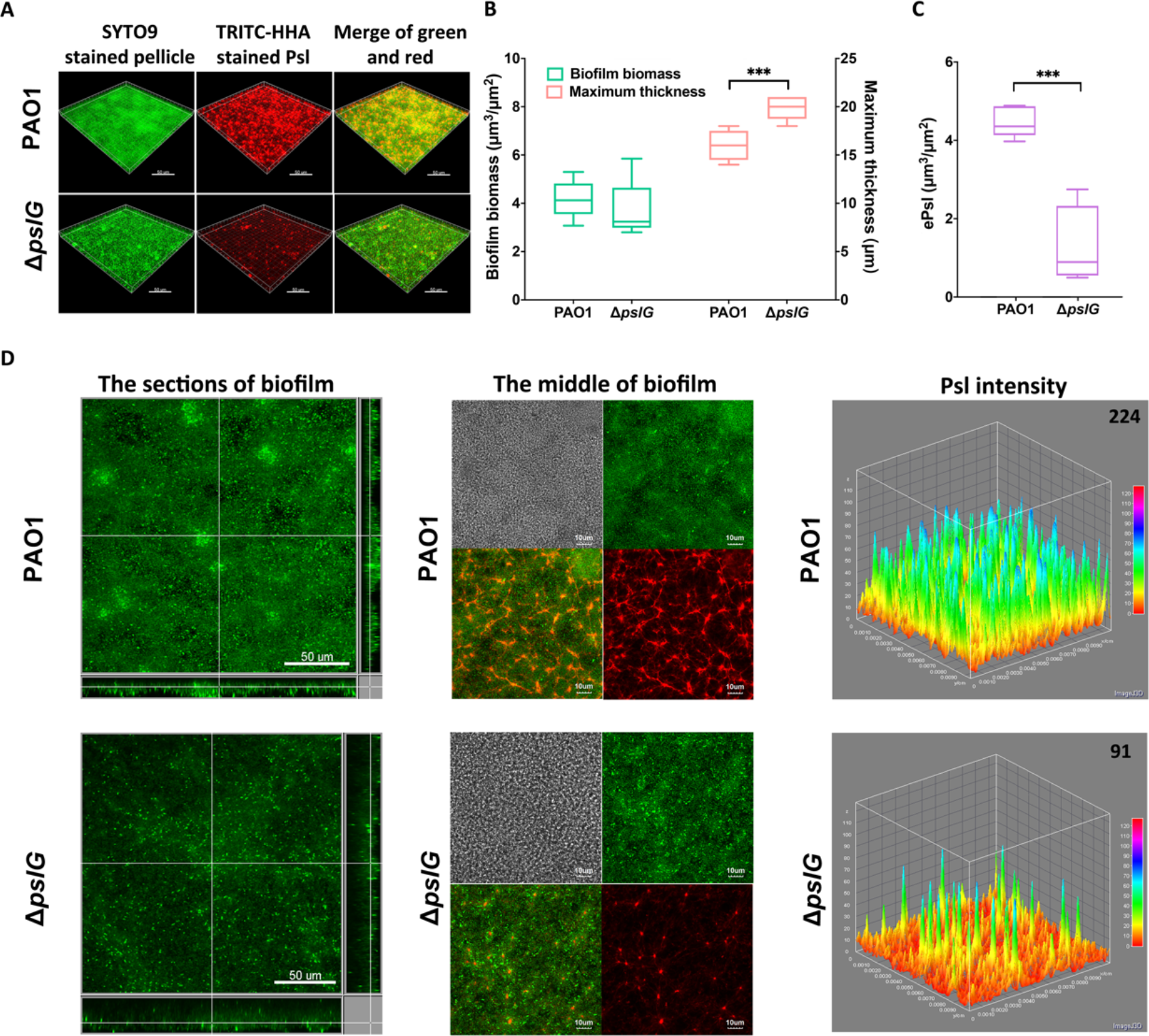
Comparison of pellicles formed by PAO1 and Δ*pslG* mutant. A: Three-dimensional images of 24-h air-liquid interface biofilms (pellicles) formed by PAO1 and Δ*pslG*. B: Biofilm biomass and maximum thickness of PAO1 or Δ*pslG* strain. C: ePsl in pellicles of PAO1 and Δ*pslG*. D: Typical section images of pellicles formed by PAO1 and Δ*pslG*. Left panel, section images showed the top-down view (square) and side view (rectangle) of corresponding pellicles. Middle panel, section images at the middle of corresponding pellicles. The distribution of bacteria (green), the fibre-like ePsl matrix (red) and corresponding DIC images (grey) were shown. Right panel, ePsl fluorescence intensity in corresponding biofilm images shown in the middle panel (the average intensity of ePsl in per μm^3^ biofilm is shown in the upper right corner). Green, SYTO9 stained bacteria, Red, TRITC-HHA stained ePsl. Statistical significances were measured using student’s t-test (***, P<0.001 when compared to PAO1). Scale bar: 50μm for A and the left panel in D; 10μm for the middle panel in D. **Figure 2-source data 1. Figure 2B and 2C source data.**

### Loss of PslG promotes the formation of microcolonies in flow-cell systems

To further understand the effect of Δ*pslG*, by employing bacterial tracking techniques, we observed Δ*pslG* cell behavior at the single-cell level. Figure 3A shows the surface coverage obtained by all tracked bacterial trajectories for a specific time period during microcolony formation. Red color indicates the surface area that has been visited by bacteria, while black color indicates a “fresh” surface area that has never been visited. Bacterial cells are shown in blue. When compared Δ*pslG* and PAO1 under the same total bacterial visits (marked as *N* in Figure 3A), the difference in the surface coverage between two strains is not obvious at *N*∼10,000. As *N* increases, the surface coverage of Δ*pslG* is clearly less than that of PAO1 (Figure 3A). At *N*∼100,000, Δ*pslG* has a surface coverage of 52 ± 6% while PAO1 has 81 ± 10%. Compared with PAO1, the less efficiency in covering the surface leads to a more hierarchical bacterial visit distribution for Δ*pslG* (that is, having a broader range of visit frequencies) (Figure 3B). Correspondingly, by fitting the distribution of bacterial visits with a power law, different power law exponents, -2.9 ± 0.1 for PAO1 and -2.5 ± 0.1 for Δ*pslG*, were obtained (Figure 3C). Such differences in bacterial visits distribution resulted that the time required to observe a visible microcolony (defined as clusters of more than 30 cells in this study, marked by dash lines in Figure 3D) in the field of view was shorter for Δ*pslG* (5.9 ± 1.7 hrs) than that for PAO1 (8.7 ± 2.4 hrs). In addition, after 10 hrs of growth, the microcolonies formed by Δ*pslG* were more compact than those formed by PAO1 (indicating by strong fluorescence intensity, Figure 3E). Taken together, these results indicate that loss of *pslG* promotes microcolonies formation by changing the surface exploration of bacteria at early stages of biofilm formation, a phenomenon that has been reported mostly for strains producing high level ePsl (Zhao *et al*., 2013).

**Figure 3.**
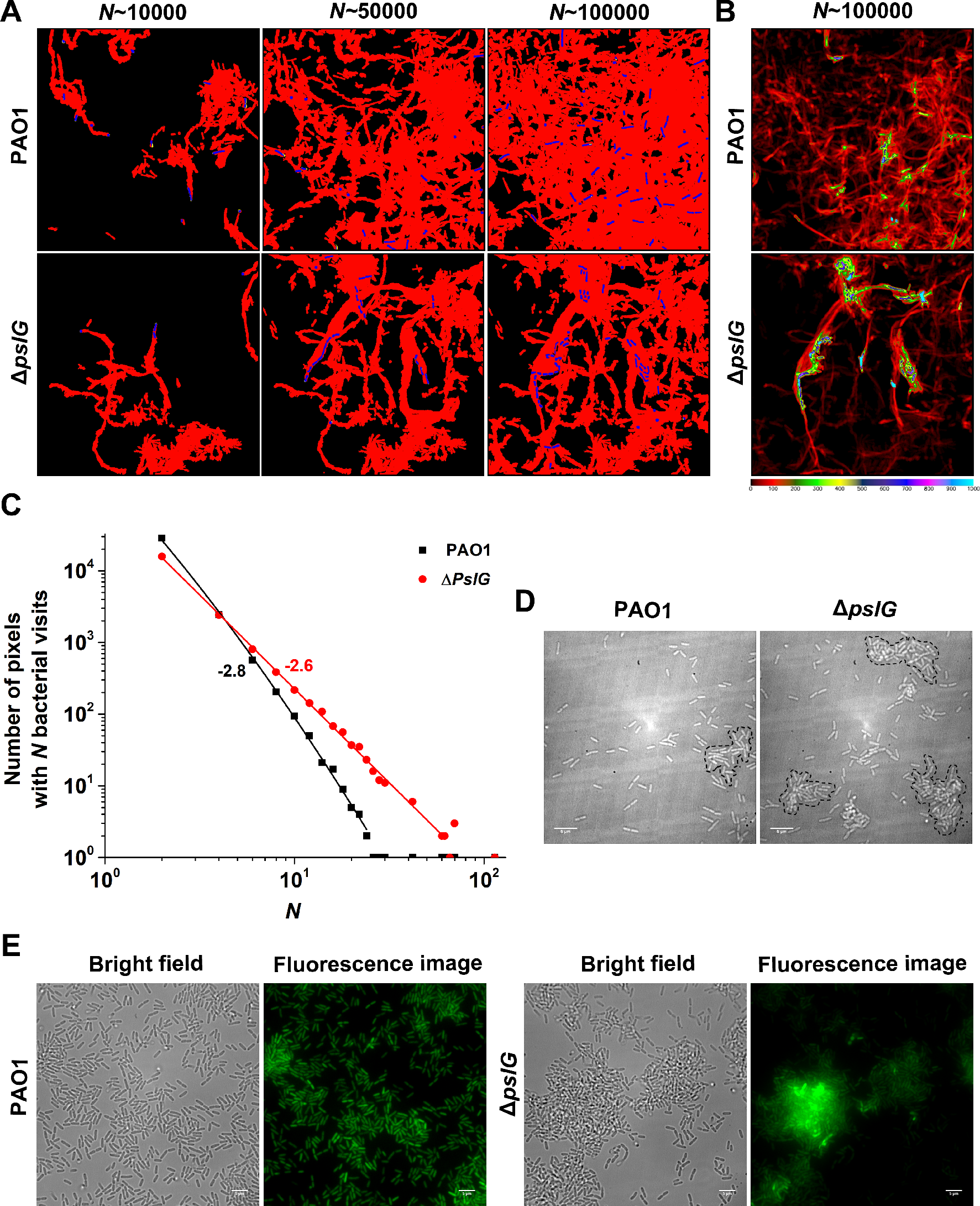
Effects of Δ*pslG* on the formation of microcolonies in flow cell systems. A: Surface coverage maps at a total of 10000, 50000 and 100000 bacterial visits for both PAO1 and Δ*pslG* cells. Red color indicates the surface area that has been visited or contaminated, while black color indicates a “fresh” surface area. Bacteria in the current frame are shown in blue. B: The intensity map of bacterial visits at *N* ∼ 100,000. The color scale black to blue represents bacterial visits of 0 to 1000. C: The graph displays the visit frequency distributions for PAO1 and Δ*pslG* of bacterial visits at *N* ∼ 100,000. D: Examples of microcolonies (enclosed by dash lines) formed by PAO1 and Δ*pslG* cells, respectively, cultured in a flow cell for about 8 hrs. E: Snapshots taken at 10 hrs after inoculation in a flow cell, showing the microcolonies formed by Δ*pslG* are more compact than those formed by PAO1. Scale bar, 5 µm. **Figure 3-source data 1. Figure 3c source data.**

### T4P-driven motility is not the main factor in promoting the microcolony formation in flow-cell systems

T4P-driven motilities on surface, such as walking and crawling, affects microcolonies formation in flow-cell systems (Conrad *et al*., 2011). To further investigate why Δ*pslG* promotes microcolonies formation, we calculated the twitching speed of bacterial cells. To minimize the effect due to possible different production of ePsl, we compared the measurements between PAO1 and WFPA801Δ*pslG* under 0.5% arabinose, under which both strains show relatively similar production of ePsl (Figure 1). The results show a slightly reduced average speed and a higher crawling percentage of WFPA801Δ*pslG* cells compared with PAO1 (Figure 1-Figure supplement 1A, B). But such differences are not statistically significant (p=0.29), indicating that the twitching motility may not be the main factor in promoting the microcolony formation.

### Lacking of *pslG* has effects on the fate and c-di-GMP distribution of daughter cells during cell division

C-di-GMP is a critical intracellular signal molecule that affects a variety of cell activities including cell motility, cell fate after division and biofilm formation. We then monitored the c-di-GMP level of cells for each cell division event by employing p*cdrA::gfp* as a reporter (Irie *et al*., 2012). Cells with a higher level of c-di-GMP show a stronger fluorescence intensity as previously described (Irie *et al*., 2012). We focused on the first bacterial division events after cells attached on the surface and analyzed the fluorescence intensity (corresponding to c-di-GMP levels) in two daughter cells right after division (Figure 4A). By comparing the fluorescence intensity of each daughter cell relative to its mother cell, the division events can be classified into three types: none of daughter cells becomes bright (none-bright), one daughter cell becomes bright (one-bright) and both of daughter cells become bright (two-bright). The results show that none-bright type is observed most frequently in both PAO1 and WFPA801Δ*pslG* strains, which has an occurrence probability of ∼60% for PAO1 and ∼52% for WFPA801Δ*pslG*. However, interestingly, WFPA801Δ*pslG* shows a relatively higher probability of two-bright type (∼17%) than that of PAO1 (∼11%) (Figure 4B and Figure 4-Figure supplement 1). Thus, compared with PAO1, WFPA801Δ*pslG* cells would have a higher probability to have both daughter cells with high c-di-GMP levels, which might enhance their stay on the surface, reduce their surface motility, and promote the microcolony formation.

**Figure 4.**
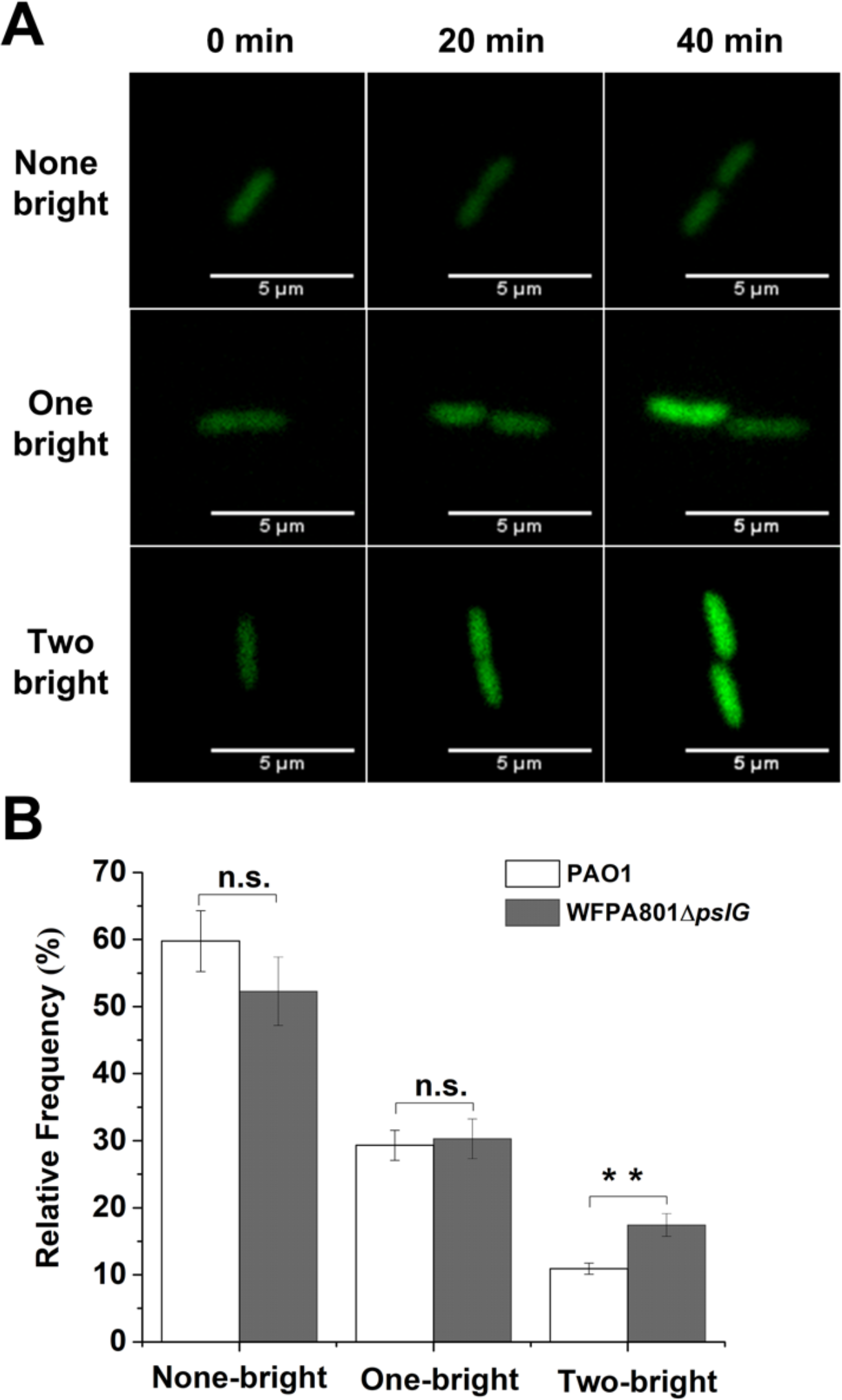
Lacking of *pslG* has effects on the fate and c-di-GMP distribution of daughter cells. A: Three types of cell division based on fluorescence intensity changes of daughter cells relative to that of their mother cell: none of daughter cells becomes bright (none-bright), one daughter cell becomes bright (one-bright), and both daughter cells become bright (two-bright). Examples given are PAO1 cells. The fluorescence is from p*cdrA*::*gfp*, which acts as a reporter for the c-di-GMP level of cells. B: The measured probability of three types of division in PAO1 and WFPA801Δ*pslG*. The total number of analyzed division events from more than three repeats is n=174 for PAO1 and n=109 for WFPA801Δ*pslG*. Statistical significances are measured using one-way ANOVA. n.s., not significant; **p < 0. 01. Scale bar, 5 µm **Figure 4-source data 1. Figure 4B source data.** **Figure 4-Figure supplement 1. The ratio of fluorescence intensity γ (γ = Idau/Imot ) of each daughter cell for each division event.** Each division event is represented as a set of two dots connected with a vertical segment, in which each dot represents one daughter cell. The horizontal dashed lines indicate the threshold value determined by 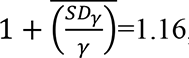, so when a daughter cell has a γ>1.16, it is considered to be brighter than its mother cell. The data are grouped based on the three types of division defined in the main text: None-bright (brown), One-bright (cyan), Two-bright (pink).

### ePsl produced by Δ*pslG* mutant has a stronger signaling function and altered structure

Irie et al. have reported that ePsl has a signaling function by stimulating intracellular c-di-GMP production (Irie *et al*., 2012). To see whether ePsl produced from Δ*pslG* has similar signaling functions, we set up a co-culture system to test the signaling functions of ePsl. The system contained an ePsl donor strain and a reporter strain. PAO1 harboring p*cdrA*::*gfp* in a plasmid was utilized as an intracellular c-di-GMP reporter strain while PAO1, WFPA800, and Δ*psl*G strains were ePsl donors respectively, in which WFPA800 was used as a negative control because it does not produce ePsl. As shown in Figure 5A, PAO1 that produces wild type level of ePsl can induce a stronger GFP fluorescence signal in the reporter stain compared to WFPA800 (Figure 5A and 5B). Surprisingly, ePsl of Δ*psl*G strain stimulated a higher fluorescence signal than that of PAO1 (Figure 5B) although Δ*psl*G mutant produces much less ePsl (about 30% of PAO1 level, Figure 1A), suggesting that ePsl synthesized from Δ*psl*G mutant has stronger signaling effect on stimulating the Aintracellular c-di-GMP production than that of wild type.

**Figure 5.**
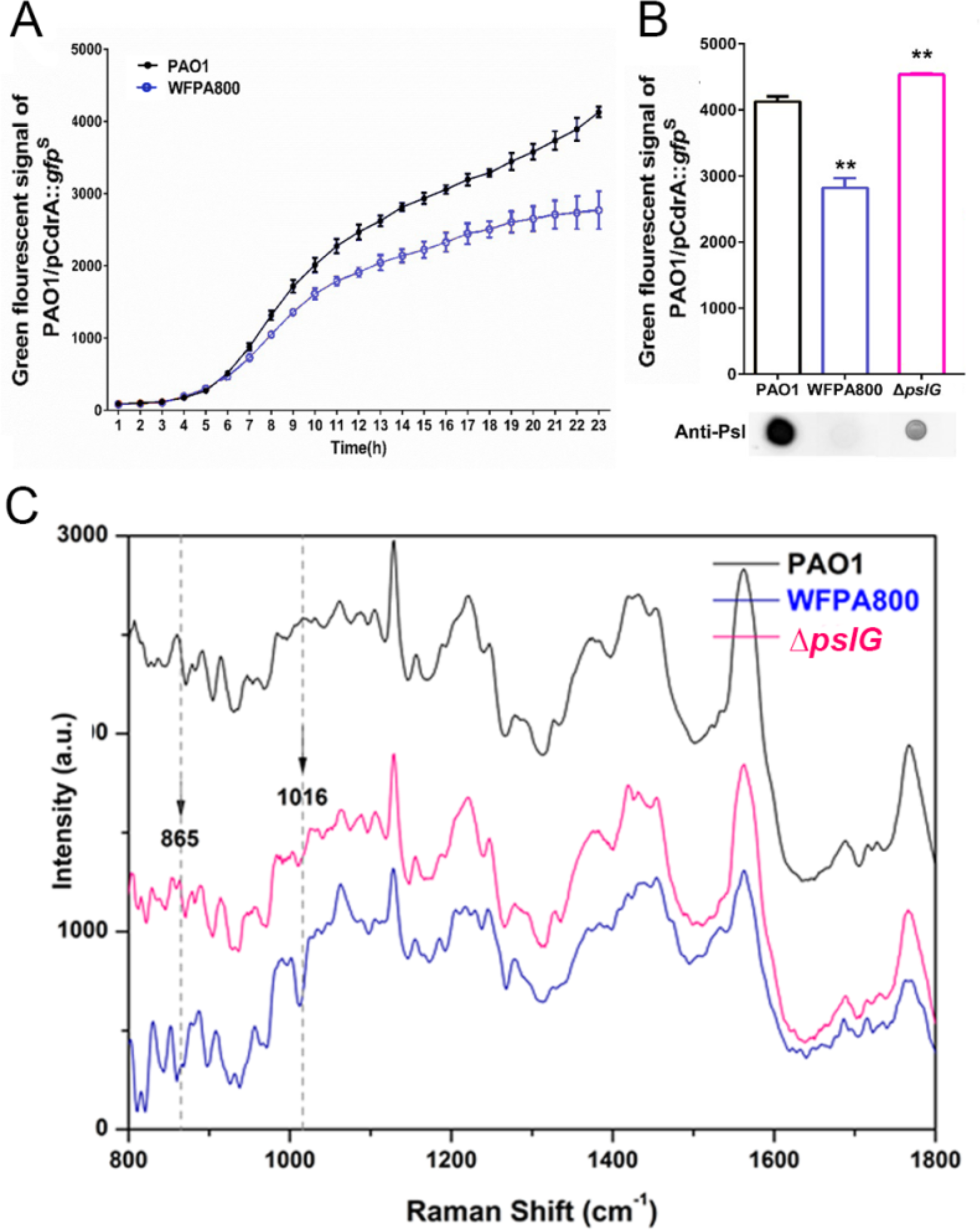
ePsl produced by *pslG* knock-out mutant has a stronger signaling function and altered structure. (A) A coculture system using p*cdrA*::*gfp* as a reporter plasmid to evaluate the intracellular c-di-GMP. The reporter strain (PAO1/p*cdrA::gfp*) was mixed with ePsl provider strain (PAO1 or a Psl-negative mutant, WFPA800) at a ratio of 1: 1, and the GFP fluorescence value in each co-culture system was recorded once per hour for 24 hours. (B) The fluorescence intensity of PAO1/p*cdrA::gfp* after 24 hours co-culture with ePsl provider strain PAO1, WFPA800 (*psl* mutant), or Δ*pslG*. ePsl production of each donor strain was shown under corresponding column. Statistical significances were measured using student’s t-test (**, p < 0. 01 when compared to PAO1). (C) Raman spectra of exopolysaccharides in biofilms of PAO1, WFPA800, and Δ*pslG*. **Figure 5-source data. Figure 5A and 5B source data.**

We then applied Raman spectroscopy to detect exopolysaccharides in the biofilm formed by PAO1 and Δ*pslG* mutant. The Δ*pslG* mutant shows a different Raman spectra at 865 cm^-1^ compared to that of PAO1 and WFPA800, indicating a change at C-C stretching and C-O-C 1,4-glycosidic link (Figure 5C). This result suggests that the structure of ePsl produced by Δ*pslG* mutant strain is different from that of PAO1, which would presumably contribute to the enhanced ePsl signaling function and the formation of microcolonies.

### Δ*pslG* shapes the localization of ePsl on bacterial periphery, leading to long chains of bacterial cells that are connected by ePsl

During bacterial tracking in flow cells, we frequently observed long chains of bacterial cells in the Δ*pslG* mutant in PAO1 background or WFPA801Δ*pslG* with arabinose induction (Figure 6A), which typically started to appear 1∼2 hours after inoculation of bacteria into a flow cell under tested conditions and could reach to about 50% of cell population at later time (Figure 6B). Such long chains of cells were not observed in strains that have intact *pslG*, such as PAO1 and WFPA801 with or without arabinose addition (Figure 6-Figure supplement 1). In a typical cell division process, two daughter cells will be disconnected from each other when the formation of septum completed. However, in *pslG* deletion mutants, the two daughter cells could not separate into physically disconnected progenies, leading to a cell chain (See one example of WFPA801Δ*pslG* in Supplementary movie S1). The length of chains varied and with chains of 4 cells were observed most frequently in either Δ*pslG* or WFPA801Δ*pslG* strain (Figure 6C). These bacterial cell chains can grow as cells continue to divide, yet some can also be broken by bending of chains, suggesting that the bacterial chains were not connected by septum. In addition, bacterial cell chains were also observed in liquid culture of Δ*pslG* mutants (data not shown), indicating that bacterial adhering on surfaces is not required for the formation of bacterial cell chains.

**Figure 6.**
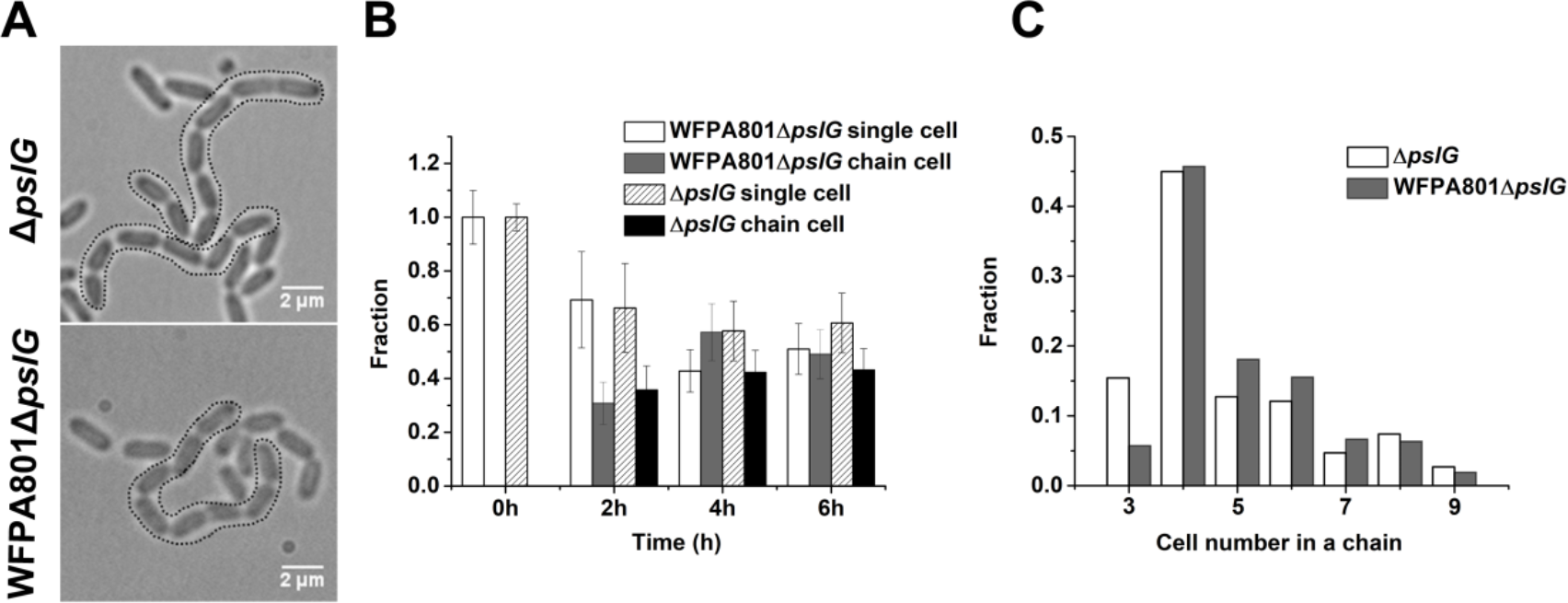
Characterization of long bacterial chains of Δ*pslG* strains. A: Examples of long bacterial chains (indicated by black dotted outlines) formed by Δ*pslG* and WFPA801Δ*pslG* cells. B: The faction of single isolated bacterial cell and cell in chains at different time points after inoculation in a flow cell. The number of analyzed picture in each strain is n=88 (about 3200 cells) for WFPA801Δ*pslG* and n=87 (about 3500 cells) for Δ*pslG*. C: The number distribution of cells consisted in a chain. The number of analyzed cells is n=301 for WFPA801Δ*pslG* and n=322 for Δ*pslG*. Scale bar, 2 µm. **Figure 6-source data 1. Figure 6B source data.** **Figure 6-source data 2. Figure 6C source data.** **Figure 6-Figure supplement 1. Examples of PAO1, Δ*pslG*, WFPA801 and WFPA801Δ*pslG* cells grown under 0% arabinose and 0.5% arabinose.** Under both conditions, long bacterial chains are observed in Δ*pslG,* but not in PAO1 and WFPA801; while in WFPA801Δ*pslG*, long chains of cells are observed under 0.5% arabinose but not under 0% arabinose. Black arrows indicate the chains of bacterial cells.

To investigate whether ePsl has any contribution on the formation of such long bacterial chains, we stained ePsl in the biofilm formed in flow cells by FITC-HHA (green fluorescence dyes FITC labeled lectin HHA). The ePsl of Δ*pslG* strains is tightly associated with bacteria compared to PAO1 and WFPA801 with arabinose (Figure 7A and Figure 7>-Figure supplement 1). We also used cell membrane stain FM4-64 to help locate cell periphery and septum. Strikingly, strong ePsl signal is often found around septa in WFPA801Δ*pslG* strain, which barely observed in WFPA801 under the same growth condition (Figure 7A).

**Figure 7.**
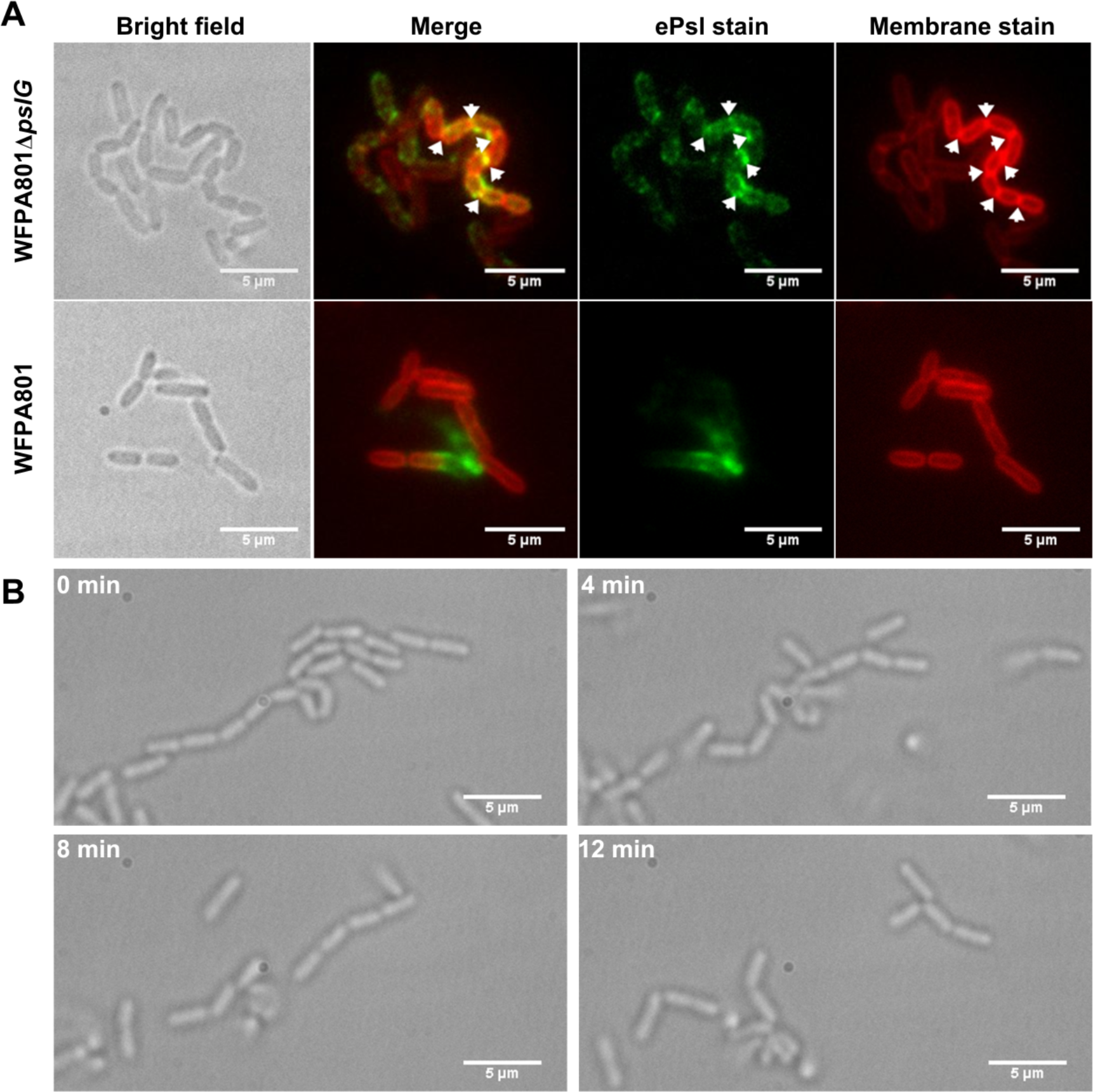
Cells in bacterial chains were connected by ePsl and can be disassembled by PslG supplied exogenously. A: Fluorescence staining of long bacterial chains formed by WFPA801Δ*pslG* cells. Staining of control samples of WFPA801 is also shown. Green shows ePsl stained by FITC-HHA, and red shows the bacterial cell membranes stained by FM4-64. Arrows in the ePsl stain image indicate the bright spotted ePsl locations, and arrows in the membrane stain image show the septum locations. B: Time–lapse images showed the break-up process of long bacterial chains of WFPA801Δ*pslG* when PslG was supplied. Scale bar, 5 µm. **Figure 7-Figure supplement 1.** An example of fluorescence staining of PAO1 cells. Green shows ePsl stained by FITC-HHA, and red shows the plasma membranes stained by FM4-64.

From the ePsl staining results, we speculate that ePsl around septa of cells might help the formation of long bacterial chains. To test this hypothesis, we first tested whether the WFPA801Δ*pslG* could form long bacterial chains under a culture condition without arabinose, under which the transcription of *psl* operon is not induced and there is no ePsl production. As shown in Figure 6-Figure supplement 1, no bacterial chains were observed in samples of WFPA801Δ*pslG* at such a no-inducer condition, suggesting the involvement of ePsl in the formation of long bacterial chains. Next, we treated the long bacterial chains with purified PslG, which has been shown to be able to disaggregate microcolonies and matured biofilms by hydrolyzing ePsl (Yu *et al*., 2015). At 4 minutes after addition of exogenous PslG, the long chains of bacterial cells were seen clearly to start to be broken up, and they were completely disconnected into single cells after 12 minutes of PslG treatment (Figure 7B) (Supplementary movie S2). Similar phenomena were also observed in the samples of Δ*pslG* (data not shown). These results together with the fact that ePsl is a “sticky” polysaccharide and PslG hydrolyzes ePsl, suggest that bacterial cells are frequently connected by ePsl in Δ*pslG* mutant strains, leading to the long bacterial cell chains. In addition, this data also suggest that the long bacterial chains might be another contributor for Δ*pslG* strains to promote microcolony formation.

Taken together, our results showed that lacking of intracellular PslG altered the structure of ePsl, shaped the localization of ePsl on bacterial periphery, enhanced the signaling function of ePsl, affected the c-di-GMP distribution in daughter cells during cell division, led to daughter cell connected together after cell division and thus the formation of bacterial chains. Consequently, Δ*pslG* exhibited fast-forming and compact biofilm microcolonies in flow cell systems and uneven bacterial distribution within a biofilm (Figure 8). Our data revealed the important roles of an intracellular glycosyl hydrolase co-transcribed with polysaccharide synthesis genes, on bacterial physiology and the biofilm development.

**Figure 8.**
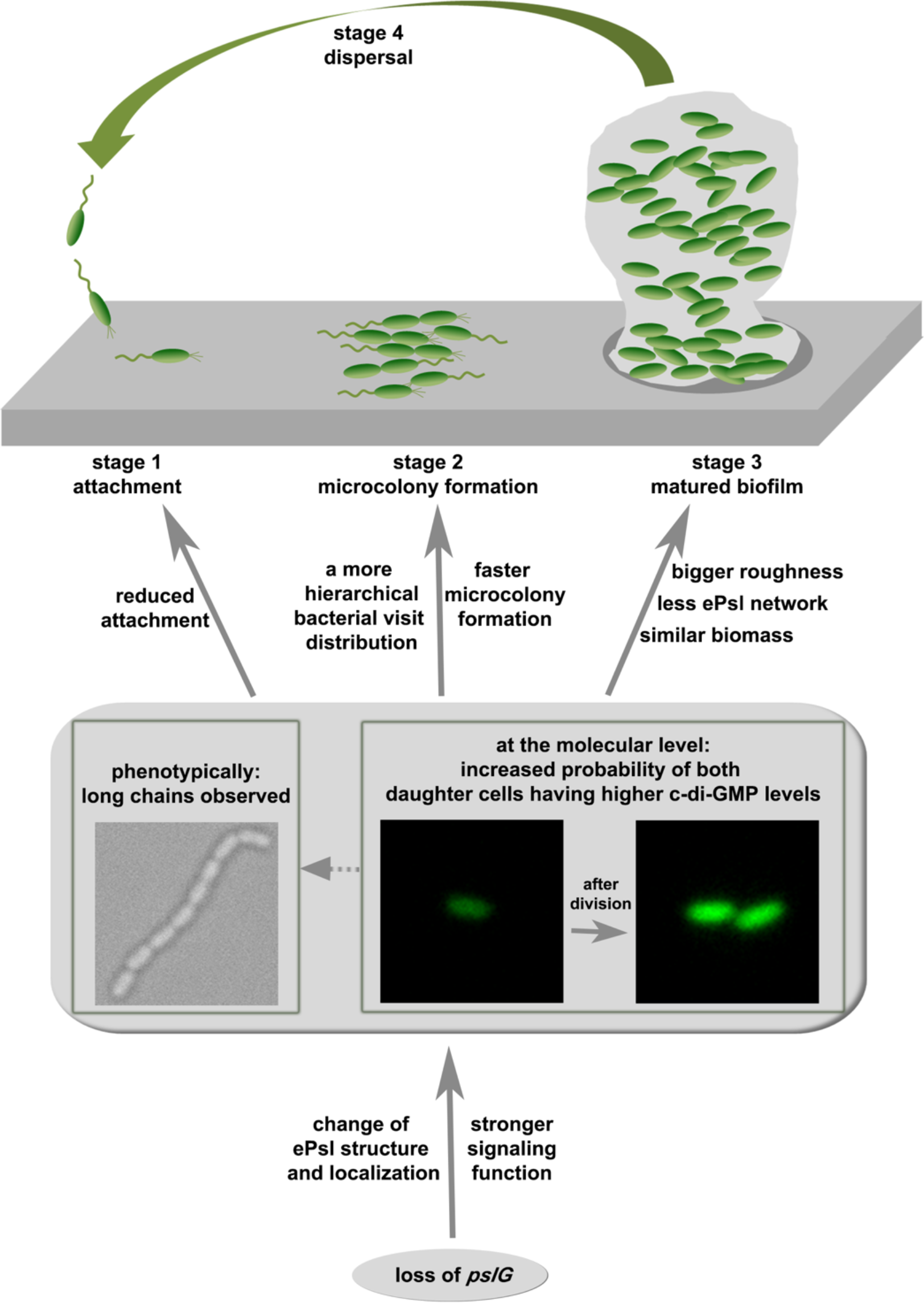
The effect of *pslG* in the biofilm development of *P. aeruginosa*.

## Discussion

In this work, by systematically studying the *pslG* knock-out mutants both at a single cell and community level, the effect of *pslG* on the bacterial physiology and surface behavior has been illustrated comprehensively. Based on our results, we propose a model for the role of *pslG* in the biofilm development of *P. aeruginosa* as the following (Figure 8).

The structural analysis of PslG has shown that it has a domain for hydrolysis of ePsl, and is typically considered to modify the Psl molecules before they are secreted out of the cell (Yu *et al*., 2015; Baker *et al*., 2015). Thus, loss of *pslG* causes a structural change in ePsl, which is evidenced by the Raman spectra (Figure 5C), and this structural change very likely alters (at least partially) the properties and functions of ePsl, such as the signaling and its localization. The consequence of this change has two aspects. One is phenotypically long chains of cells are formed. The long chains are not observed in PAO1 and WFPA801, both of which have intact *pslG*. Such long chains seem not induced by the adherence on a surface as they can also be observed in liquid cultures. Rather, there are multiple lines of evidences to support that ePsl is the main factor for the formation of long chains. Firstly, by adding PslG externally into cell cultures, long chains were observed to disassemble into single cells, presumably due to the hydrolysis of ePsl by PslG as shown in literature (Yu *et al*., 2015; Baker *et al*., 2015; Zhang *et al*., 2018). Secondly, such long chains are not observed in WFPA801Δ*pslG* under 0% arabinose (i.e., strains with no ePsl production). Thirdly, the fluorescence staining experiments also show that ePsl are often localized around septa of cells, where daughter cells are supposed to be disconnected after division.

The other aspect as a consequence of the change in ePsl due to the loss of *pslG* is, at the molecular level, the probability of both daughter cells having a higher level of c-di-GMP than that of their mother cell in a division event is increased. The increased level of c-di-GMP then would result in reduced cell motility and promote cells to transit to biofilm style (Romling *et al*., 2013). This may also help cells to form long chains by reducing the breakage of chains due to reduced cell motility and increased ePsl production.

As it is widely known that ePsl plays a very important role in the biofilms of *P. aeruginosa*, then both aforementioned aspects would have an effect on the biofilm development at different stages. At the attachment stage, reduced swimming motility, which can be caused by the higher level of c-di-GMP, can contribute to the reduced surface attachment observed on microtiter surfaces. Long chains of cells may also contribute to the reduced swimming motility. However, under our experimental setup, we cannot measure the swimming of long chains in liquid cultures when they move toward to the air-liquid interface. After cells attached, both the formation of long chains and the increased level of c-di-GMP in two daughter cells will help both daughter cells to stay on the surface, and result in a more hierarchical bacterial visit distribution (Figure 3), which then lead to an earlier formation of microcolony. As cells continue to grow and proliferate, such differences in cell behavior between WFPA801Δ*pslG* and PAO1 finally result in matured biofilms with different structures as shown in pellicles formed at the air-liquid interface, where pellicles of WFPA801Δ*pslG* are rougher and have less web-like Psl structures than those of PAO1 although their biomasses are similar. We note that the aforementioned effects of *pslG* on the biofilm development is also dependent on ePsl, because if there is no ePsl production, there will be no much difference between with and without *pslG*.

PslG has been shown to be an efficient ePsl degrader *in vitro* and in biofilm matrix, yet its roles within a bacterium is not completely understood. Wu et al have shown that the glycoside hydrolytic activity of PslG is important for ePsl production (Wu *et al*., 2019). Baker et al. have proposed that PslG might be responsible for the degradation of Psl polysaccharide stock in the periplasmic space in *P. aeruginosa* (Baker *et al*., 2015). Beyond the aforementioned roles of PslG, our observations on the bacterial behavior resulted from the loss of *pslG* further expand our understandings of PslG functions. To the best of our knowledge, the ability of a glycosyl hydrolase to affect bacterial phenotype and c-di-GMP levels has not been reported until this work. It would be also very interesting to see in the future whether other glycosyl hydrolases in different bacterial species can have similar functions.

In summary, in this study, we have provided a comprehensive analysis on the effect of *pslG* on the biofilm development of *P. aeruginosa*. Our results indicate that although *pslG* is not essential for synthesis of ePsl, it plays an important role in regulating the proper functions of ePsl, and loss of *pslG* results in malfunction of ePsl, which then cause changes in both morphology and surface behavior of bacterial cells through ePsl-mediated interactions. This work shed light on better understanding the role of PslG and would be helpful in developing new ways for biofilm control through *pslG*-based ePsl regulation.

## Materials and Methods

### Bacterial strains and growth conditions

All *P. aeruginosa* stains used in study were listed in table S1. *P. aeruginosa* stains were grown at 37 °C in LB without sodium chloride (LBNS) or Jensen’s, a chemically defined media (Jensen *et al*., 1980). Biofilms of *P. aeruginosa* were grown in Jensen’s medium at 30 °C. L-arabinose (Sigma) was used as inducer for genes transcribed from P_BAD_ promoter in *P. aeruginosa*. Antibiotics for *P. aeruginosa* were added at the following concentrations: gentamicin 30 µg/ml; ampicillin 100 µg/ml; carbenicillin 300 µg/ml. For *Pseudomonas* selection media, irgasan at 25 µg/ml was used.

The *psl*-inducible strains WFPA801*ΔpslG* was constructed in accordance with WFPA801 (the promoter of *psl* operon in PAO1 was replaced by *araC*-P_BAD_-*psl*) (Ma *et al*., 2006). Briefly, plasmid pMA9 (Ma *et al*., 2006) was transferred into *pslG* deletion mutant by conjugation (Wu *et al*., 2019). For single recombination mutant selection, LBNS plates with 30 µg/ml gentamycin and 25 µg/ml irgasan were used; for double recombination mutant selection, LBNS plates containing 10% sucrose were used.

### Bacterial attachment on microtiter dish

The assay was done as described previously with modifications (Ma *et al*., 2006; O’Toole, 2011). Overnight culture was 1/100 diluted into Jensen’s media (with or without arabinose) and incubated at 37 °C with shaking until the OD_600_ reached 0.5. The 100μl of such culture was inoculated into 96-well PVC microtiter dish (BD Falcon), and incubated at 30 °C for 30 min. Then the planktonic and loosely adherent bacteria cells were washed off by rinsing the plate in water. The remaining surface-attached cells were stained by 0.1% crystal violet, solubilized in 30% acetic acid, and finally measured the value of OD_560_.

### Swimming motility assay

Swimming motility assay was performed as preciously described (Zhao *et al*., 2018). Briefly, strains were grown overnight on LBNS plates. Single colony was stab-inoculated with a sterile toothpick on the surface of LBNS plates (0.3% BD Bacto Agar). Plates were incubated upright at 37 °C overnight. Swimming zones were measured accordingly.

### ePsl dot-blotting

Strains were incubated in Jensen’s medium with shaking at 30°C for 24 h. Cells of an OD600 of 10 were collected by centrifugation to extract crude bacterial surface-bound polysaccharides. Pellet was resuspended in 100 µl of 0.5 M EDTA, and boiled at 100°C for 5 min. After centrifugation at 13,000 g for 10 min, the supernatant fraction was treated with 0.5 mg/ml proteinase K at 60°C for 1 h and proteinase K was then inactivated at 80°C for 30 min. ePsl immunoblotting was performed as previously described using ePsl antibody (Byrd *et al*., 2009). ImageJ software was used to quantify the immunoblot data. The protein concentration of each sample culture was measured by a BCA protein assay kit (Thermo) to ensure the same amount of cell lysate was used in each experiment.

### Flow cell assembly, sterilization and washing of the system

Flow cells made of polycarbonate were purchased from the Department of Systems Biology, Technical University of Denmark. Each flow cell has three identical rectangle channels (40 × 4 × 1 mm^3^) and was assembled by attaching a cover glass as substratum as previously described (Sternberg and Tolker-Nielsen, 2006). The assembled flow cell was connected to a syringe through a 0.22 µm filter (Millipore) using silicon tubing. Then the whole system was sterilized overnight with 3% H_2_O_2_ at 3 ml/h using a syringe pump (Harvard Apparatus). After sterilization, autoclaved, deionized water was used to wash the whole system overnight. Before inoculation of bacteria into the flow cell, the system was flushed for 5 minutes at a flow rate of 30ml/h by Jensen’s medium using a syringe pump (Harvard Apparatus). Then the medium flow was stopped and 1ml of a diluted bacteria culture (OD_600nm_ ∼ 0.01) were injected directly into the channel of the flow cell using a 1ml syringe equipped with a needle. A 5-minute incubation period was allowed after inoculation to let cells attaching to the surface, which was then followed by a medium flow with a large flow rate of 30 ml/h for 5 minutes to wash out floating cells. After that the flow rate was set to 3 ml/h, and image recording was started. In this work, the flow cell experiments were conducted at 30 °.

### Biofilms and image acquisition

Pellicles (air-liquid interface biofilms) were grown in glass chambers (Chambered # 1.5 German Coverglass System, Nunc) with a glass coverslip at the bottom of each chamber as described previously (Wang *et al*., 2013). 1/100 dilution of a saturated (overnight) culture in Jensen’s media for *P. aeruginosa* was inoculated into the chamber, and incubated at 30 °C for 24h. The ePsl was stained with lectin TRITC-HHA (EY lab, INC) at 100 µg/ml for 2 h in the dark. Then bacteria were strained with SYTO9 (5 µM final concentration, Molecular Probes, Invitrogen) for 15 min. Fluorescent images were obtained using a FV1000 CLSM (Olympus, Japan). The excitation/emission parameters for TRITC-HHA and SYTO9 were 554 nm/570 nm and 480 nm/500 nm, respectively. CLSM-captured images were analyzed using a COMSTAT software (Heydorn *et al*., 2000).

For flow cell experiments, the flow was stopped before staining. ePsl was stained with lectin FITC-HHA (EY lab, INC) at 100 µg/ml for 20 minutes in the dark, then the flow was running for a short time to flush out the non-binding dye. Subsequently, bacteria were tagged by Gfp or stained by cell membrane stain FM4-64 (10 µM final concentration, Molecular Probes) for 2 mins in the dark (without flow). Next, the flow was resumed to flush out the dye and ready for examination under microscope. Images were captured using an EMCCD camera (Andor iXon) on a Leica DMi8 microscope equipped with Zero Drift autofocus system. The image size is 66.5 µm × 66.5 µm (1,024×1,024 pixels). The images were recorded with a 100× oil objective (plus 2× magnifier).

### Examination of *P. aeruginosa* biofilms by Raman spectroscopy

*P. aeruginosa* pellicles (air-liquid biofilms) were grown in Jensen’s medium in 24 well plate (NEST biotech Co., Ltd.) at 30 °C for 24 h. Then pellicles were transferred into the quartz tank for Raman spectrum detection. Raman spectra were acquired with an integration time of 60 s and a laser power of 25 mW at 633 nm under Acton spectroscopy 2300 Raman microscope system. Originpro 8.1 software was applied for data processing and analysis.

### Detection of ePsl signaling function on stimulating c-di-GMP using co-culture system

The c-di-GMP levels were determined using p*cdrA::gfp* as a reporter as described previously (Byrd *et al*., 2009). The growth curve and green fluorescent signal of PAO1/p*cdrA::gfp* were measured via recording the OD600 values and the corresponding GFP fluorescence (Ex/Em 488/520) by a Synergy H4 hybrid reader (BioTek).The promoter activity was calibrated as the relative fluorescence divided by the OD600. In co-culture system, PAO1 harboring plasmid p*cdrA*::*gfp* was the reporter strain to indicate the level of intracellular c-di-GMP. PAO1, WFPA800, and Δ*psl*G strains were ePsl donor strains, respectively.

### Single cell tracking image analysis

Images were processed and analyzed in the same way as described in reference (Zhang *et al*., 2018). Simply, 16-bit greyscale images were first converted to binary images for the detection of bacteria with a standard image processing algorithm. Geometry information of cells such as center position, size and aspect ratio etc. were then collected. Bacterial trajectories were obtained by connecting cell positions in all frames of a time series, from which bacterial motion can be measured and analyzed.

For quantitatively comparing the fluorescence intensity of cells containing p*cdrA*::*gfp* reporter between mother cell and daughter cells, first the fluorescence intensity of each cell *I* was measured by the averaged fluorescence intensity value within the area enclosed by the cell envelope. The mother cell was measured when it irreversibly attached to the surface (typically 40∼50 minutes before the division completed), and the daughter cells were measured right after the division (i.e., the two daughter cells are completely separated. In practice, the daughter cells were measured within 10 minutes right after the division completed due to the 10-mimute time interval for the fluorescent image recording). Then the ratio of fluorescence intensity γ between each daughter cell (*I*_dau_) and its mother cell (*I*_mot_) was calculated, γ = *I*_dau_/*I*_mot_. The relative standard deviation was estimated by 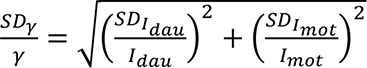 Here, 𝑆𝐷_$_ refers to standard deviation of γ. Similar for 𝑆𝐷_*’()_ and 𝑆𝐷_*-./_. We define a daughter cell to be fluorescent brighter than its mother cell if 𝛾 > 1 + 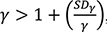 here 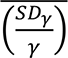 is the averaged value of the relative standard deviation for all analyzed division events.

A cluster is an aggregation of multiple cells. We used a minimum distance criterion to judge whether a cell belonged to a cluster or not. If the minimum distance between any point of the scrutinized cell body and any point of any cell body of the cluster, is smaller than 0.5µm (i.e., about one width of abacterial cell), then the scrutinized cell is considered to belong to the cluster. Otherwise not.

## Acknowledgements

This work is supported by the National Key R&D Program of China (2018YFA0902102, 2019YFC804104, and 2019YFA0905501) and the National Natural Science Foundation of China (91951204, 21621004, 32070033). The funders had no role in the study design, data collection and interpretation, or the decision to submit the work for publication.

## Data availability

Data presented in this study are available from the corresponding author upon reasonable request.

## Movie Legends

**Supplementary Movie S1.** An example movie of the formation of WFPA801Δ*pslG* long cell-chain. The movie was taken at a frame interval of 5 min for 3.5h and was played back at 5 fps. Scale bar, 5 µm.

**Supplementary Movie S2.** An example movie of the degradation of WFPA801Δ*pslG* long cell-chain. The movie was taken at a frame interval of 1 min for 0.5h and was played back at 5 fps. Scale bar, 5 µm.

## Key resources table

| Reagent type (species) or resource | Designation | Source or reference | Identifiers |
| --- | --- | --- | --- |
| Strain, strain background (*Pseudomonas aeruginosa*) | PAO1 | Holloway, 1955 | Prototroph |
| Strain, strain background (*Pseudomonas aeruginosa*) | $\Delta pslG$ | Wu *et al.*, 2019 | In-frame deletion of *pslG* |
| Strain, strain background (*Pseudomonas aeruginosa*) | WFPA800 | Ma *et al.*, 2006 | ePsl-negative, the *psl* operon promoter deletion mutant, $\Delta P_{psl}$ |
| Strain, strain background (*Pseudomonas aeruginosa*) | WFPA801 | Ma *et al.*, 2006 | ePsl-inducible strain, the promoter of *psl* operon is replaced by *araC*-P<sub>BAD</sub>, $\Delta P_{psl}$ / P<sub>BAD</sub>-*psl* |
| Strain, strain background (*Pseudomonas aeruginosa*) | WFPA801$\Delta pslG$ | This study | In-frame deletion of *pslG* in WFPA801 background |
| Recombinant DNA reagent | *pcdrA::gfp* | Rybtke *et al.*, 2012 | GFP reporter of promoter *cdrA* activity, to indicate intracellular c-di-GMP level. Derived from pUCP22Not-RNase III-gfp(ASV)-T0-T1, Amp<sup>R</sup>, Gm<sup>R</sup> |
| Antibody (anti-ePsl) |  | Byrd *et al.*, 2009 | Exopolysaccharide Psl specific antibody |
| Chemical compound, dye | TRITC-HHA | EY-lab, INC | Fluorescent labeled lectin HHA |
| Chemical compound, dye | FITC-HHA | EY-lab, INC | Fluorescent labeled lectin HHA |
| Chemical compound, dye | Syto9 | Invitrogen, Molecular probes | Green-fluorescent nucleic acid stain |
| Chemical compound, dye | FM4-64 | Invitrogen, Molecular probes | Lipophilic Styryl Dye. |

---

**Figure 1-Figure supplement 1.**
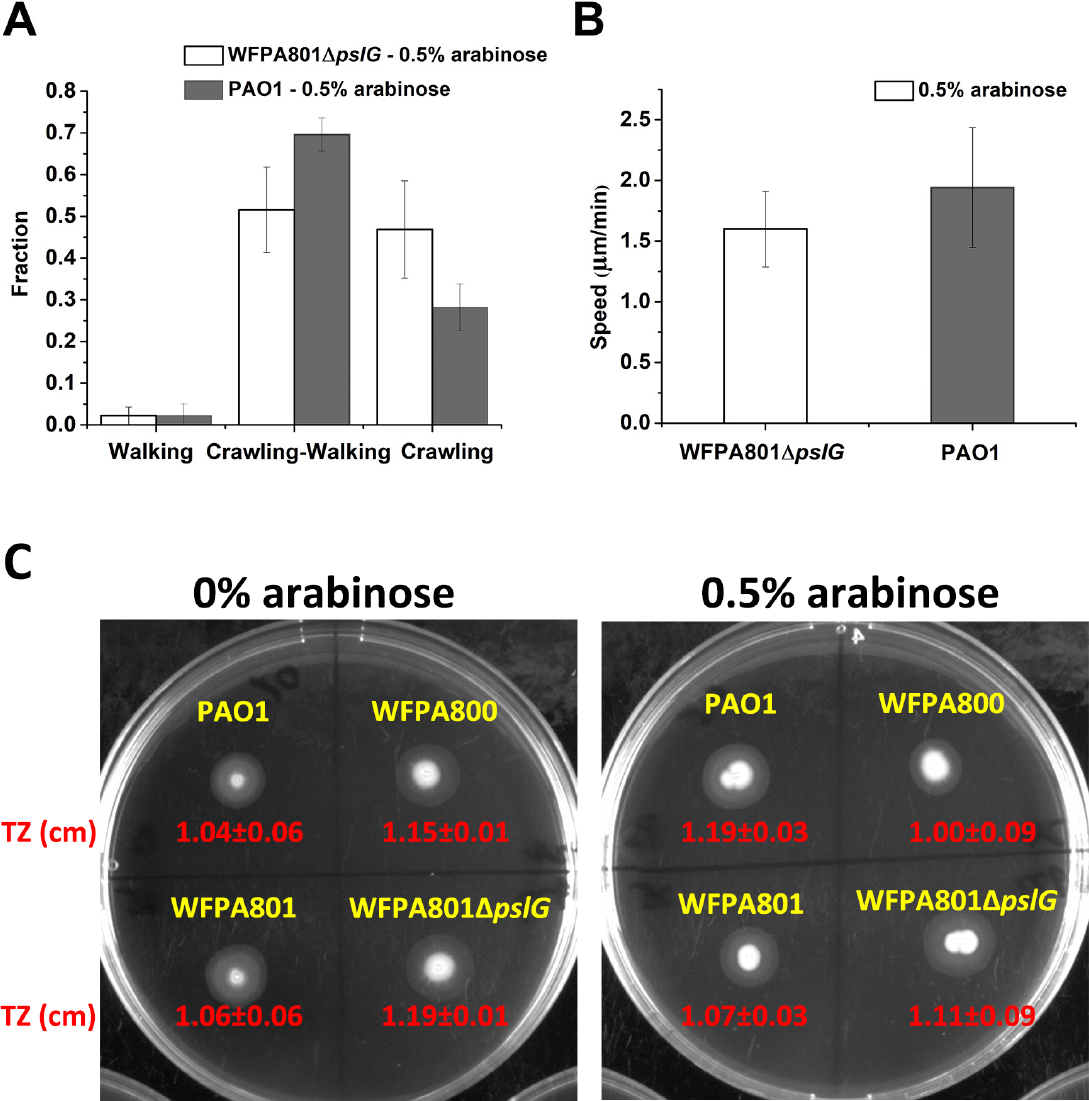
The influence of Δ *pslG* on T4P-driven twitching motility. A: Tracking T4P-driven bacterial walking and crawling on the glass surfaces of flow-cell systems. Shown are the fractions of cells that crawl only, walk only and both crawl and walk for WFPA801Δ*pslG* and PAO1. B: Speed of cell twitching for WFPA801Δ*pslG* and PAO1. The number of analyzed cells is n=496 for PAO1 and n=576 for WFPA801Δ*pslG*. C: The effect of Δ*pslG* on the twitching zone.

**Figure 4-Figure supplement 1.**
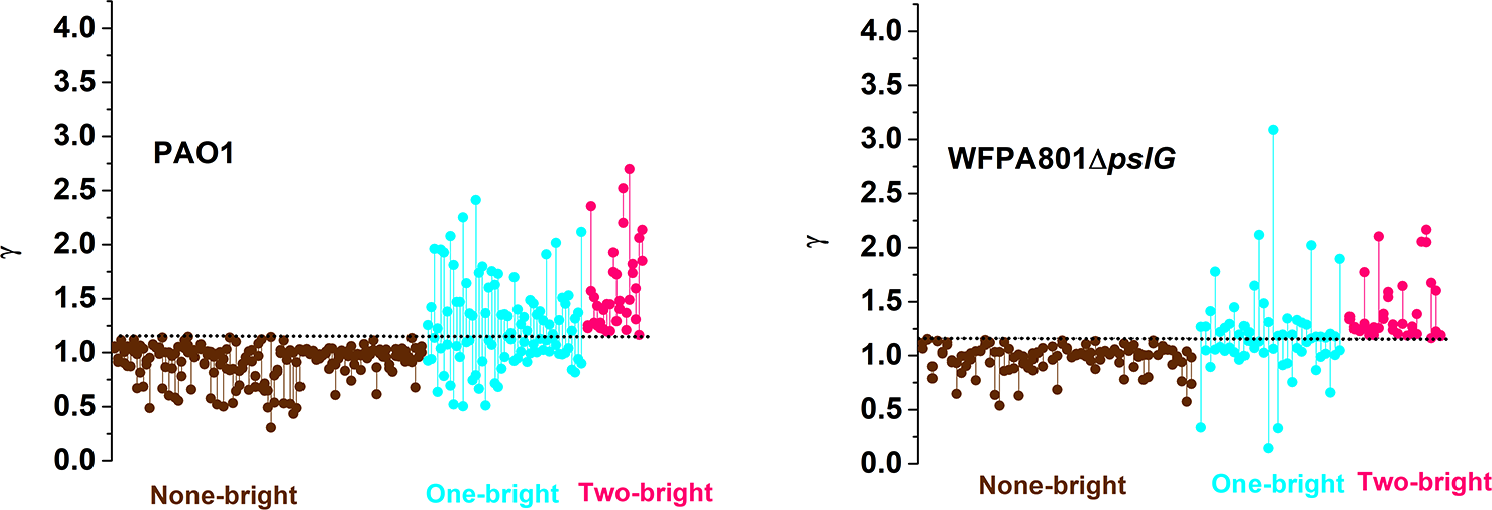
The ratio of fluorescence intensity γ (γ = *I*_dau_/*I*_mot_ ) of each daughter cell for each division event. Each division event is represented as a set of two dots connected with a vertical segment, in which each dot represents one daughter cell. The horizontal dashed lines indicate the threshold value determined by 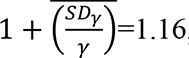, so when a daughter cell has a γ>1.16, it is considered to be brighter than its mother cell. The data are grouped based on the three types of division defined in the main text: None-bright (brown), One-bright (cyan), Two-bright (pink).

**Figure 6-Figure supplement 1.**
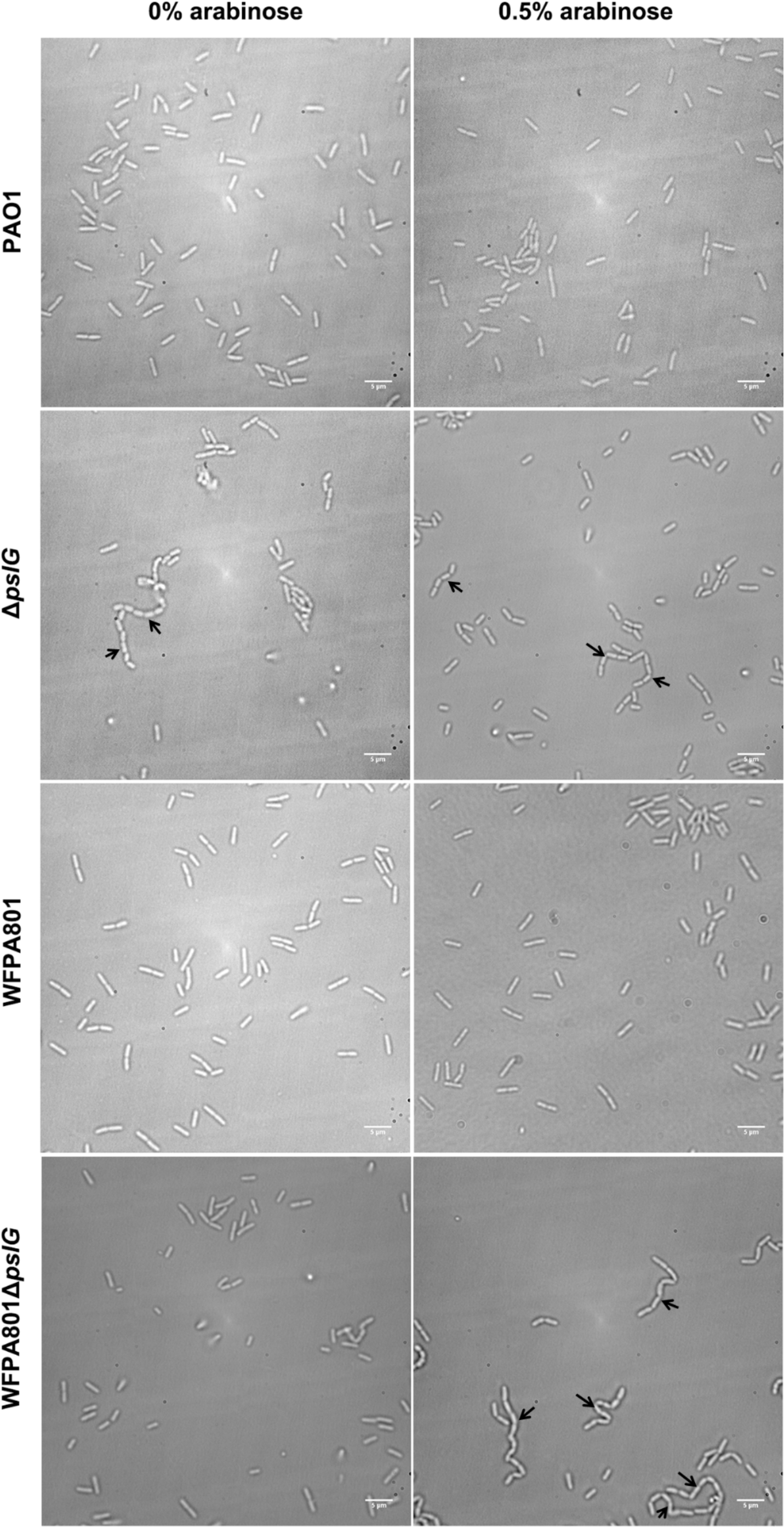
Examples of PAO1, Δ*pslG*, WFPA801 and WFPA801Δ*pslG* cells grown under 0% arabinose and 0.5% arabinose. Under both conditions, long bacterial chains are observed in Δ*pslG,* but not in PAO1 and WFPA801; while in WFPA801Δ*pslG*, long chains of cells are observed under 0.5% arabinose but not under 0% arabinose. Black arrows indicate the chains of bacterial cells.

**Figure 7-Figure supplement 1.**
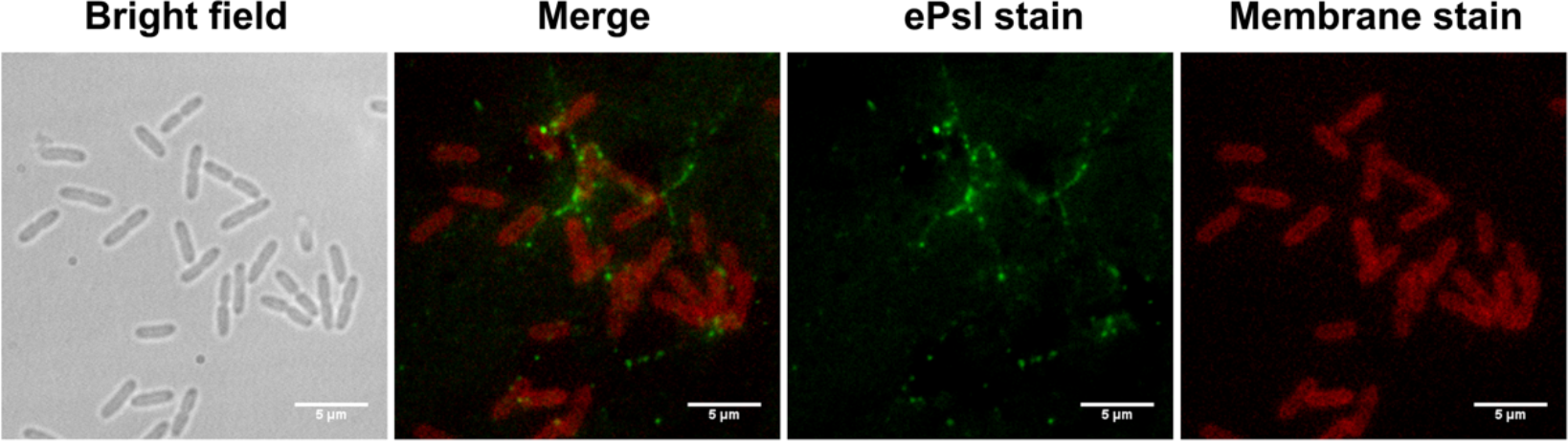
An example of fluorescence staining of PAO1 cells. Green shows ePsl stained by FITC-HHA, and red shows the plasma membranes stained by FM4-64.

## References

Baker P, Whitfield GB, Hill PJ, Little DJ, Pestrak MJ, Robinson H, Wozniak DJ, Howell PL. 2015. Characterization of the *Pseudomonas aeruginosa* glycoside hydrolase PslG reveals that its levels are critical for Psl polysaccharide biosynthesis and biofilm formation. Journal of Biological Chemistry 290:28374–28387. DOI: 10.1074/jbc.M115.674929, PMID: 26424791

Billings N, Millan M, Caldara M, Rusconi R, Tarasova Y, Stocker R, Ribbeck K. 2013. The extracellular matrix Component Psl provides fast-acting antibiotic defense in *Pseudomonas aeruginosa* biofilms. PLoS Pathog 9:e1003526. DOI: 10.1371/journal.ppat.1003526, PMID: 23950711

Bruzaud J, Tarrade J, Coudreuse A, Canette A, Herry JM, Taffin de Givenchy E, Darmanin T, Guittard F, Guilbaud M, Bellon-Fontaine MN. 2015. Flagella but not type IV pili are involved in the initial adhesion of *Pseudomonas aeruginosa* PAO1 to hydrophobic or superhydrophobic surfaces. Colloids and Surfaces B:Biointerfaces 131:59–66. DOI: 10.1016/j.colsurfb.2015.04.036, PMID: 25950497

Byrd MS, Sadovskaya I, Vinogradov E, Lu H, Sprinkle AB, Richardson SH, Ma L, Ralston B, Parsek MR, Anderson EM, Lam JS, Wozniak DJ. 2009. Genetic and biochemical analyses of the *Pseudomonas aeruginosa* Psl exopolysaccharide reveal overlapping roles for polysaccharide synthesis enzymes in Psl and LPS production. Molecular Microbiology 73:622–638. DOI: 10.1111/j.1365-2958.2009.06795.x, PMID: 19659934

Christen M, Kulasekara HD, Christen B, Kulasekara BR, Hoffman LR, Miller SL. 2010. Asymmetrical distribution of the second messenger c-di-GMP upon bacterial cell division. Science 328:1295–1297. DOI: 10.1126/science.1188658, PMID: 20522779

Colvin KM, Irie Y, Tart CS, Urbano R, Whitney JC, Ryder C, Howell PL, Wozniak DJ, Parsek MR. 2012. The Pel and Psl polysaccharides provide *Pseudomonas aeruginosa* structural redundancy within the biofilm matrix. Environmental Microbiology 14:1913–1928. DOI: 10.1111/j.1462-2920.2011.02657.x, PMID: 22176658

Conrad, JC., Gibiansky, ML., Jin, F., Gordon, VD., Motto, DA., Mathewson, MA., Stopka, WG., Zelasko, DC., Shrout, JD., Wong, GCL. 2011. Flagella and pili-mediated near-surface single-cell motility mechanisms in *P. aeruginosa*. Biophysical Journal 100:1608–1616. DOI: 10.1016/j.bpj.2011.02.020, PMID: 21463573

Costerton JW, Lewandowski Z, Caldwell DE, Korber DR, Lappinscott HM. 1995. Microbial biofilms. Annual Review of Microbiology 49:711–745. DOI: 10.1146/annurev.mi.49.100195.003431, PMID: 8561477

Flemming HC, Wingender J. 2010. The biofilm matrix. Nature Reviews Microbiology 8:623–633. DOI: 10.1038/nrmicro2415, PMID: 20676145

Franklin MJ, Nivens DE, Weadge JT, Howell PL. 2011. Biosynthesis of the *Pseudomonas aeruginosa* extracellular polysaccharides, alginate, Pel, and Psl. Frontiers in Microbiology 2:167. DOI: 10.3389/fmicb.2011.00167, PMID: 21991261

Govan JRW, Deretic V. 1996. Microbial pathogenesis in cystic fibrosis: mucoid *Pseudomonas aeruginosa* and *Burkholderia cepacia*. Microbiology and Molecular Biology Reviews 60:539–574. DOI: 10.1128/mr.60.3.539-574.1996, PMID: 8840786

Haussler S, Parsek MR. 2010. Biofilms 2009: new perspectives at the heart of surface-associated microbial communities. Journal of Bacteriology 192:2941–2949. DOI: 10.1128/JB.00332-10, PMID: 20382760

Heydorn A, Nielsen AT, Hentzer M, Sternberg C, Molin S. 2000. Quantification of biofilm structures by the novel computer program COMSTAT. *Microbiology (Reading*, England) 146 (Pt 10):2395–2407. DOI: 10.1099/00221287-146-10-2395, PMID: 11021916

Holloway BW. 1955. Genetic recombination in *Pseudomonas aeruginosa*. Journal of General and applied Microbiology 13:572–581. DOI: 10.1099/00221287-13-3-572, PMID: 13278508

Irie Y, Borlee BR, O’Connor JR, Hill PJ, Harwood CS, Wozniak DJ, Parsek MR. 2012. Self-produced exopolysaccharide is a signal that stimulates biofilm formation in *Pseudomonas aeruginosa*. PNAS 109: 20632–20636. DOI: 10.1073/pnas.1217993109, PMID: 23175784

Jensen SE, Fecycz IT, and Campbell JN. 1980. Nutritional factors controlling exocellular protease production by *Pseudomonas aeruginosa*. Journal of Bacteriology 144:844–847. DOI: 10.1128/jb.144.2.844-847.1980, PMID: 6776099

Laventie BJ, Sangermani M, Estermann F, Manfredi P, Planes R, Hug I, Jaeger T, Meunier E, Broz P, Jenal U. 2019. A surface-induced asymmetric program promotes tissue colonization by *Pseudomonas aeruginosa*. Cell Host & Microbe 25:140–152. DOI: 10.1016/j.chom.2018.11.008, PMID: 30581112

Lyczak JB, Cannon CL, Pier GB. 2000. Establishment of *Pseudomonas aeruginosa* infection: Lessons from a versatile opportunist. Microbes and Infection 2:1051–1060. DOI: 10.1016/s1286-4579(00)01259-4, PMID: 10967285

Ma L, Jackson KD, Landry RM, Parsek MR, Wozniak DJ. 2006. Analysis of *Pseudomonas aeruginosa* conditional psl variants reveals roles for the psl polysaccharide in adhesion and maintaining biofilm structure postattachment. Journal of Bacteriology 188:8213–8221. DOI: 10.1128/JB.01202-06, PMID: 16980452

Ma L, Conover M, Lu H, Parsek MR, Bayles K, Wozniak DJ. 2009. Assembly and development of the *Pseudomonas aeruginosa* biofilm matrix. PLoS Pathogens 5:e1000354. DOI: 10.1371/journal.ppat.1000354, PMID: 19325879

Mishra M, Byrd MS, Sergeant S, Azad AK, Parsek MR, McPhail L, Schlesinger LS, Wozniak DJ. 2012. *Pseudomonas aeruginosa* Psl polysaccharide reduces neutrophil phagocytosis and the oxidative response by limiting complement-mediated opsonization. Cellular Microbiology 14:95–106. DOI: 10.1111/j.1462-5822.2011.01704.x, PMID: 21951860

O’Toole, G.A., and Kolter, R. 1998. Flagellar and twitching motility are necessary for *Pseudomonas aeruginosa* biofilm development. Molecular Microbiology 30: 295–304. DOI: 10.1046/j.1365-2958.1998.01062.x, PMID: 9791175

O’Toole GA. 2011. Microtiter dish biofilm formation assay. Jove-Journal of Visualized Experiments 30:2437. DOI: 10.3791/2437, PMID: 21307833

Ramsey DM, Wozniak DJ. 2005. Understanding the control of *Pseudomonas aeruginosa* alginate synthesis and the prospects for management of chronic infections in cystic fibrosis. Molecular Microbiology 56:309–322. DOI: 10.1111/j.1365-2958.2005.04552.x, PMID: 15813726

Romling U, Galperin MY, Gomelsky M. 2013. Cyclic di-GMP: the first 25 years of a universal bacterial second messenger. Microbiology and Molecular Biology Reviews 77:1–52. DOI: 10.1128/MMBR.00043-12, PMID: 23471616

Rybtke MT, Borlee BR, Murakami K, Irie Y, Hentzer M, Nielsen TE. 2012. Fluorescence-based reporter for gauging cyclic di-GMP levels in *Pseudomonas aeruginosa*. Applied and Environmental Microbiology 78: 5060–5069.

Sternberg C, Tolker-Nielsen T. 2006. Growing and analyzing biofilms in flow cells. Current Protocols in Microbiology Chapter 1:Unit 1B.2. DOI: 10.1002/9780471729259.mc01b02s00, PMID: 18770573

Stewart, PS, Costerton, JW. 2001. Antibiotic resistance of bacteria in biofilms. The Lancet Child & Adolescent Health 358:135–138. DOI: 10.1016/s0140-6736(01)05321-1, PMID: 11463434

Stoodley P, Sauer K, Davies DG, Costerton JW. 2002. Biofilms as complex differentiated communities. Annual Review of Microbiology 56:187–209. DOI: 10.1146/annurev.micro.56.012302.160705, PMID: 12142477

Tseng BS, Zhang W, Harrison JJ, Quach TP, Song JL, Penterman J, Singh PK, Chopp DL, Packman AI, Parsek MR. 2013. The extracellular matrix protects *Pseudomonas aeruginosa* biofilms by limiting the penetration of tobramycin. Environmental Microbiology 15:2865–2878. DOI: 10.1111/1462-2920.12155, PMID: 23751003

Wang S, Parsek MR, Wozniak DJ, Ma LZ. 2013. A spider web strategy of type IV pili-mediated migration to build a fibre-like Psl polysaccharide matrix in *Pseudomonas aeruginosa* biofilms. Environmental Microbiology 15:2238–2253. DOI: 10.1111/1462-2920.12095, PMID: 23425591

Wu H, Wang D, Tang M, Ma LZ. 2019. The advance of assembly of exopolysaccharide Psl biosynthesis machinery in *Pseudomonas aeruginosa*. MicrobiologyOpen 8:e857. DOI: 10.1002/mbo3.857, PMID: 31070012

Yu S, Su T, Wu H, Liu S, Wang D, Zhao T, Jin Z, Du W, Zhu MJ, Chua SL, Yang L, Zhu D, Gu L, Ma LZ. 2015. PslG, a self-produced glycosyl hydrolase, triggers biofilm disassembly by disrupting exopolysaccharide matrix. Cell Research 25:1352–1367. DOI: 10.1038/cr.2015.129, PMID: 26611635

Zhang J, He J, Zhai C, Ma LZ, Gu L, Zhao K. 2018. Effects of PslG on the surface movement of *Pseudomonas aeruginosa*. Applied and environmental microbiology 84:e00219–18. DOI: 10.1128/AEM.00219-18, PMID: 29728385

Zhao K, Tseng BS, Beckerman B, Jin F, Gibiansky ML, Harrison JJ, Luijten E, Parsek MR, Wong GC. 2013. Psl trails guide exploration and microcolony formation in *Pseudomonas aeruginosa* biofilms. Nature 497:388–391. DOI: 10.1038/nature12155, PMID: 23657259

Zhao T, Zhang Y, Wu H, Wang D, Chen Y, Zhu M, Ma LZ. 2018. Extracellular aminopeptidase modulates biofilm development of *Pseudomonas aeruginosa* by affecting matrix exopolysaccharide and bacterial cell death. Environmental Microbiology Reports 10: 583–593. DOI: 10.1111/1758-2229.12682, PMID: 30047246

